# Using coverage-based rarefaction to infer non-random species distributions

**DOI:** 10.1101/2020.04.14.040402

**Authors:** Thore Engel, Shane A. Blowes, Daniel J. McGlinn, Felix May, Nicholas J. Gotelli, Brian J. McGill, Jonathan M. Chase

## Abstract

Understanding how species are non-randomly distributed in space, as well as how the resulting spatial structure of diversity responds to ecological, biogeographic and anthropogenic drivers is a critical piece of the biodiversity puzzle. However, most metrics that quantify the spatial structure of diversity (i.e., community differentiation), such as Whittaker’s classical β-diversity metric are influenced by sampling effects. As a result, these measures are influenced by species pool size, species abundance distributions and numbers of individuals. Null models have been proposed to evaluate the degree of differentiation among communities due to spatial structuring relative to that expected from sampling effects. However, to date, these null models do not accommodate the influence of sample completeness (i.e. the proportion of the species pool in the sample). Here, we develop an approach that makes use of individual- and coverage-based rarefaction and extrapolation. Using spatially explicit simulations, we show that our derived metric, *β_C_*, captures changes in intraspecific aggregation independently of changes in the species pool size. We then provide two case studies examining spatial structure in forest plots spanning latitudinal gradients: (1) a re-analysis of the “Gentry” plot dataset, and (2) comparing a high diversity plot in Barro Colorado Island, Panama with a low diversity plot in Harvard Forest, Massachusetts, USA. We find no evidence for systematic changes in spatial structure with latitude in these datasets. As it is rooted in biodiversity sampling theory and explicitly controls for sample completeness, our approach represents an important advance over existing null models for spatial aggregation. Potential applications of the approach range from better descriptors of biogeographic diversity patterns to the consolidation of local and regional diversity trends in the current biodiversity crisis.

**Open research statement:** The novel code for the calculation of *β_C_* can be found in supplementary material S4. Empirical data sets utilized for this research are as follows: Phillips & Miller (2002), Orwig et al. (2015), Condit et al. (2019). Our research repository including the novel code is also available at https://github.com/t-engel/betaC and will be uploaded to Zenodo upon acceptance of this manuscript.

## Introduction

Organisms are non-randomly distributed across the globe and understanding spatial patterns of species diversity from ecological samples remains a central challenge (Gaston, 2000; McGill, 2011a; Worm and Tittensor 2018). Spatial structure in species diversity (e.g. species turnover from site to site) is typically quantified by one or more metrics of compositional dissimilarity or spatial β-diversity (Anderson et al. 2011). Measures of β-diversity offer a mathematical link between local (i.e. α) and regional (i.e. γ) species diversity, and can, for example, shed light onto the metacommunity processes that shape biological assemblages (Chase & Myers, 2011), inform biodiversity conservation (Socolar, Gilroy, Kunin & Edwards, 2016), and help understand the provisioning of ecosystem functions and services (Mori, Isbell, & Seidl, 2018).

Although β-diversity is conceptually appealing, its quantification and interpretation is often ambiguous (Tuomisto 2010a, b; Anderson, 2011). Whittaker’s (1960) multiplicative β-diversity (γ/ α), for example, is commonly thought to represent the sort of community differentiation that arises due to nonrandom distributions of species (e.g. species turnover or intraspecific spatial aggregation). However, this and other measures of β-diversity are also influenced by the number of samples, the size of the regional species pool, the shape of the regional species abundance distribution (SAD), and the number of individuals captured by the samples (McGill, 2011, Chase & Knight, 2013, Chase et al, 2018). This makes it a challenge to compare and interpret patterns of β-diversity and related measures along biogeographic gradients (e.g. Kraft et al, 2011). To disentangle the effect of spatial aggregation from the non-spatial components that influence β-diversity (SAD or relative proportion of rare species, species pool size), empirical studies have frequently adopted null model approaches that compare the observed patterns against a null expectation that simulates spatial randomness by shuffling individuals among sites (Chase et al., 2011; Kraft et al. 2011). However, the exact formulation of the null expectation and its deviation (β-deviation) remains debated (Qian, Wang & Zhang, 2012; Kraft et al., 2012; Mori, Fujii, Kitagawa & Koide, 2015; Tucker et al., 2016; Xing & He, 2020). In particular, the null model approach has been criticized because it overlooks the influence of the completeness of the samples (Ulrich et al. 2017, Sreekar et al. 2018).

Much of the ambiguity surrounding measures of β-diversity and its null expectations can be understood in terms of sampling effects and sample completeness (i.e., the proportion of species in the species pool captured by sampling). For instance, regions with large species pools are expected to exhibit high β-diversity simply because local samples only capture a small and incomplete portion of the total diversity; this can lead to strong, but spurious differentiation among local samples (Chase et al. 2011; Kraft et al., 2011). This is not to say that this kind of sample differentiation is not meaningful, but it reflects the species pool (or the inability of local samples to sample it) rather than non-random species distributions. Similarly, sampling effects can “inflate” β-diversity when there are many rare species in an assemblage, or when the total community density is relatively low (i.e., widespread species remain undetected in most samples) (Barwell, Isaac & Kunin 2015). Although such sampling effects are ubiquitous in ecological studies (Colwell & Coddington, 1994; Gotelli & Colwell, 2001), sampling theory is not well developed with respect to β-diversity (Wolda 1981; Beck, Holloway, & Schwanghart, 2013). For example, Chao & Chiu (2016) developed a framework to unify different approaches to community differentiation, but they state clearly that their approach ignores such sampling issues. There have also been attempts to develop asymptotic β-diversity metrics (Chao, Chazdon, Colwell, & Shen, 2005) but these have been found to show strong biases when tested on simulated and empirical data (Beck Holloway, & Schwanghart, 2013; Cardoso, Borges, & Veech, 2009). While rarefaction-based approaches are commonly used to address sampling effects at a single scale by standardizing diversity to a common number of individuals (Gotell and Colwell 2001) or to equal levels of sample completeness (Chao and Jost 2012), these approaches have been rarely applied to concepts related to β-diversity (but see Olszewski 2004, Dauby and Hardy 2012, Stier et al. 2016, Chase at el, 2018).This is despite the fact that the null models used in detecting deviations from random expectations in β-diversity (e.g. Kraft et al. 2011, Xing & He, 2020) are based on largely similar concepts (i.e., difference between observed and expected measures of diversity).

In what follows, we consider non-random spatial distributions through the lens of the individual-based rarefaction curve and combine existing approaches (Chase et al. 2018, McGlinn et al. 2019) with coverage-based standardization. Specifically, we compare rarefaction curves taken from subsets of samples (i.e., an α-scale curve) to those from the entire set of samples (i.e., a γ-scale rarefaction curve), using a constant γ-scale coverage (i.e., an estimate of sample completeness). From this, we obtain a metric, which we call *β_C_*, that estimates the degree of spatial structure in the assemblage independently of the species pool size and the SAD. We emphasize that our goal here is not to develop a better measure of β-diversity per se, as it is true that Whittaker’s β-diversity and relatives have many useful properties for discerning biodiversity scaling (e.g., Jost 2007, Tuomisto 2010a,b, Chao & Chiu 2016). Instead, our goal is to develop a measure that allows us to discern the magnitude of spatial structuring within a given regional assemblage. Building on rarefaction and sampling theory has the advantage that we can evaluate sample completeness and bypass the shuffling algorithms and estimates of beta-deviation inherent to previous null-model approaches. We test our method on simulated spatial point patterns with different degrees of spatial structure (intraspecific spatial aggregation) and varying species pool sizes, and apply it to two empirical datasets to examine variation in spatial structure along a latitudinal gradient of tree diversity.

### Individual-based rarefaction and extrapolation (IBRE)

Our approach is based on individual-based rarefaction and extrapolation (IBRE), which is a common method to standardize species richness estimates (Hurlbert, 1971; Gotelli & Colwell, 2001, Chao & Jost, 2012). IBRE curves describe the non-linear scaling relationship between the number of individuals in a sample and expected species richness (i.e., rarefied richness). The shape of the curve is determined by the size of the species pool and the relative abundances of species in that pool, which is often referred to as the Species Abundance Distribution (SAD, McGill et al. 2007). The slope at any point along the curve is related to the estimated sample completeness for the number of individuals sampled at that point. For smaller than observed sample sizes, the expected number of species can be interpolated using the following formula (Hurlbert, 1971):

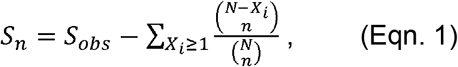

where *S_n_* is the rarefied richness, or the expected number of species for *n* individuals (n<N), *S_obs_* is the observed number of species, *N* is the observed number of individuals in the sample, and *X_i_* is the number of individuals of the *i^th^* species.

For larger than observed sample sizes, the expected number of species can be estimated using the following extrapolation formula (Chao & Jost, 2012):

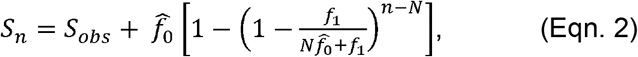

where 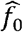 is the estimated number of unseen species, estimated as:

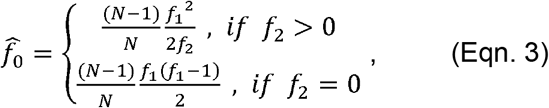

and *f_1_* and *f_2_* are the observed numbers of singletons and doubletons (i.e., species represented by one or two individuals), respectively. Extrapolation of species richness is considered unbiased, though only recommended for sample sizes up to two times the observed sample size (Chao et al, 2014).

### Spatial structure through the lens of the IBRE curve

By constructing IBRE curves from samples at two or more nested spatial scales, we can assess intraspecific spatial aggregation (Olszewski 2004, Dauby and Hardy 2012, Chase et al. 2018, McGlinn et al. 2019). Like most classical approaches to estimating diversity partitioning, we define α-diversity as the mean number of species within a given sample or subset of localized samples, and γ-diversity as the total number of species from multiple pooled samples or local subsets of samples (Tuomisto, 2010a). Accordingly, the α-scale IBRE curve is derived by calculating the IBRE curve from each individual sample and then averaging the all samples 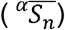 while the γ-scale curve consists of *S_n_* values calculated from the pooled sample (*^γ^S_n_*). The α-scale is influenced by turnover (i.e., spatial structure) among samples within the assemblage, whereas the γ-scale breaks up any spatial structure by randomly accumulating individuals from all samples. If species are distributed randomly among samples (i.e., there is no aggregation), the α-and γ-scale IBRE curves sit on top of each other (Fig. 1). Downward and upward deviations of the α-scale curve, then, would be interpreted as intraspecific aggregation and overdispersion, respectively (Chase et al., 2018, McGlinn et al. 2019). The γ-scale IBRE is conceptually very similar to abundance-based null expectations (e.g. Kraft et al., 2011, Xing & He, 2019), but it uses an analytical formula rather than a shuffling algorithm. Furthermore, rather than comparing the observed β-diversity to a null distribution of β-diversity, it directly compares the observed α-scale IBRE curve to the null-expectation given by the γ-scale IBRE curve.

**Figure 1:**
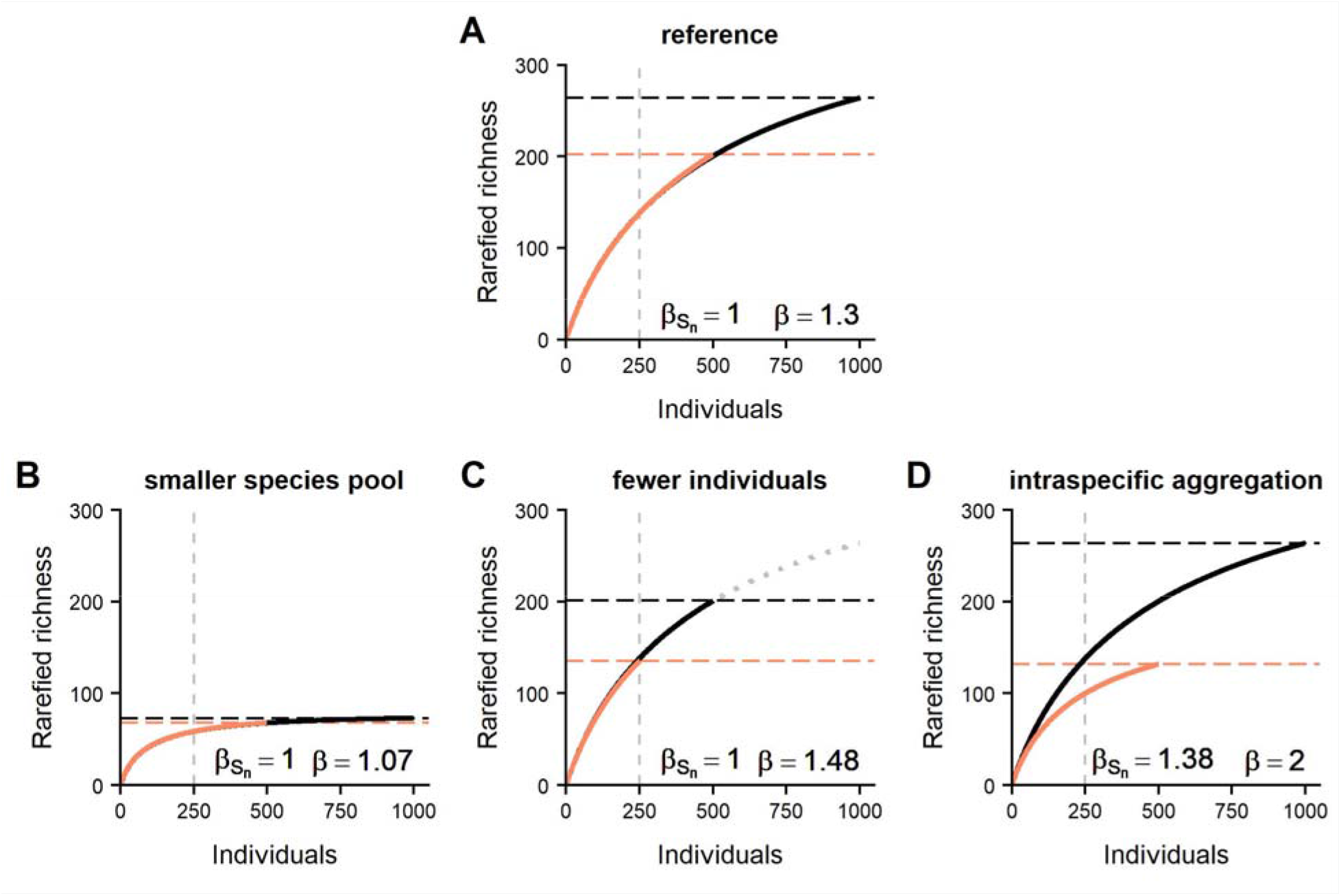
Examples of two-scale individual-based rarefaction curves for (A) a hypothetical species pool size of 450, and how they respond to, (B) reduced species pool size / altered SAD (species pool size of 80), (C) reduced numbers of individuals and (D) and changes to patterns of within species aggregation. Orange curve: α scale, black curve: γ scale. Dashed lines represent observed species richness at α and γ scale and Whittaker’s β-diversity (β) can be illustrated as the height ratio of the two. values are calculated for n = 250 individuals on all panels (dashed vertical lines). Dotted grey curve in panel C: reference curve (from A) to aid comparison.

Using IBRE curves (only interpolation shown for simplicity), Figure 1 illustrates how β-diversity of a reference assemblage (Fig. 1A) responds to changes in the size of the species pool (Fig. 1B), the numbers of individuals (Fig. 1C) and intraspecific spatial aggregation (Fig. 1D). Whittaker’s β-diversity 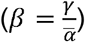 is represented as the height ratio of the two curves at the respective right-hand end of the curves (dashed horizontal lines). In each of the four examples, *β* > 1, but for very different underlying reasons. Only the assemblage underlying Fig. 1D exhibits spatial turnover in species composition (due to aggregation). In the other cases (A-C), differentiation only emerges due to a sampling effect (i.e., a ‘more-individuals’ effect between the α- and γ-scale).

Due to the nonlinear shape of the IBRE curve, the sampling effect depends on the regional SAD (Fig. 1B) and the number of individuals sampled (Fig. 1C). Chase et al. (2018) suggested that when calculated at a common number of individuals (n), the ratio of rarefied richness (S_n_) calculated between the γ and α scales, termed *β_S_n__*, could provide an indication of the degree of intraspecific aggregation, or non-randomness in the distribution of species in the assemblage, independent of any sampling effect (see also McGlinn et al. 2019). *β_S_n__* is related to the metrics developed by Olszewski (2004) and Dauby and Hardy (2012), who compared the differences of IBRE curves at their base (n=2), rather than at *n* individuals. When assemblages have a random spatial structure, *β_S_n__* is expected to equal 1 regardless of species pool and sample size (Fig 1 A-C). Conversely, *β_S_n__* values larger than 1 reflect spatial aggregation or species turnover among sites in the region (Fig 1 D).

While the deviation between α- and γ-scale IBRE curves (i.e. *β_S_n__* ≠ 1) is due to spatial structure, its magnitude is contingent on the value of *n* and the shape of the curves (i.e. the size and evenness of the species pool). Thus, as we will illustrate below, *β_S_n__* is biased when comparing the degree of aggregation among regions where species pools and shapes of the γ-scale IBRE curves change (e.g., along biogeographical gradients). To visualize this problem, consider two assemblages each composed of two patches, but which differ in the size of their regional species pool (500 vs 100 species Fig. 2A and 2B, respectively). Supposing that both assemblages have complete species turnover between their respective patches, Figure 2 shows the IBRE curves that we would expect if we sampled 500 individuals from each patch in the large (Fig. 2A) and small (Fig. 2B) species pools. Note how the γ-curve from the small species pool is much closer to its asymptote than the one from the large species pool (slope of grey tangential lines). This difference in completeness has implications for the values of *β_S_n__*. Although both assemblages are maximally and equally structured at the patch scale, the relative deviation between α and γ is substantially higher in the small species pool (indicated by *β_S_n__*). Asymptotically, both assemblages have a theoretical β-diversity of 2 (i.e. complete turnover), but at any common sample size *β_S_n__* differs between them due to the difference in sample completeness that is associated with species pool size (Fig. 2C).

**Figure 2:**
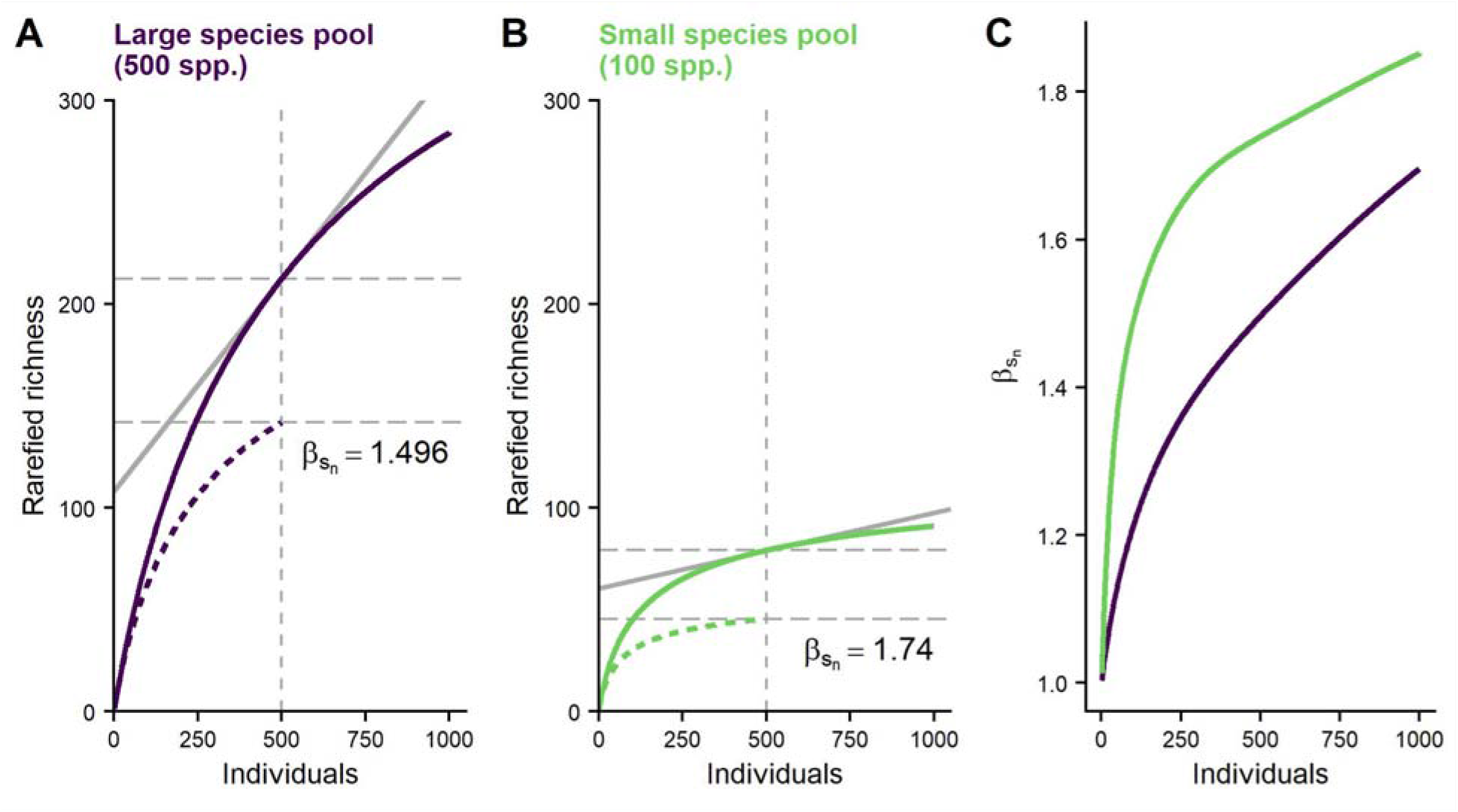
β_S_n__ is affected by species pool size in, aggregated communities. Two-scale IBRE curves for (A) a large species pool and (B) a small species pool. Solid curve: γ scale; dashed curve: α scale; dashed vertical grey line: number of individuals (n=500) used for the calculation of β_S_n__ (i.e,. the ratio of the horizontal dashed lines). Grey solid line shows the slope of the γ scale IBRE that relates to sample coverage (the steeper the slope of this line, the less complete the sample). (C) β_S_n__ plotted as a function of the corresponding sample size (n). dashed vertical line is the same as in (A) and (B).

### Coverage-based rarefaction and extrapolation

To account for the differences in species pool size and sample completeness, we extend *β_S_n__* to include the concept of coverage-based rarefaction (Chao and Jost, 2012). Sample coverage is a measure of sample completeness that ranges from 0 to 1 and refers to the “proportion of the total number of individuals in a community that belong to the species represented in the sample” (Chao & Jost, 2012). Sample coverage depends on the sample size and the species abundance distribution of the underlying assemblage. It can be estimated from the number of rare species in a sample (Good, 1953; Chao & Shen, 2010). As it is directly related to the steepness of the IBRE curve, expected coverage can also be estimated for any sample size along the curve using the following equations (Chao & Jost, 2012):

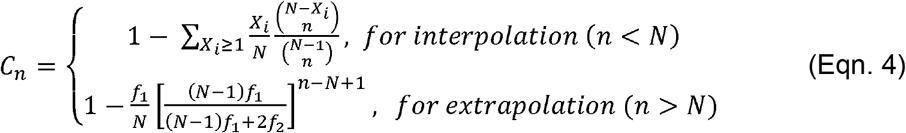

where *C_n_* is the expected coverage for a subsample of sample size *n*. *N* is the total number of individuals in the sample, *X_i_* is the number of individuals of the *i^th^* species, and *f_1_* and *f_2_* are the numbers of singletons and doubletons.

For coverage-based standardization, Eqn 4 can be solved numerically to determine how many individuals, *n*, are necessary to obtain a given target coverage, *C_target_*. This is computed by calculating *C_n_* for every possible *n* and choosing the one that minimizes the difference between *C_n_* and *C_target_* Subsequently, IBRE can be used to standardize the diversity estimate to a sample size of *n*, and thus the desired coverage level *C_target_* (e.g., Hsieh, Ma, & Chao, 2016).

### Introducing *β_C_*

By calculating *β_S_n__* for equal γ-scale coverage, we can resolve the species pool dependence when making comparisons across assemblages. Specifically, rather than keeping sample-size (*n*) constant when comparing across assemblages, we instead maintain a consistent sample coverage at the γ-scale (*C_target_*), and refer to *β_S_n__* standardized by sample coverage as *β_C_*. Figure 3 illustrates this approach using the same example with large (Fig. 3A) and small (Fig. 3B) species pools. By allowing *n* to vary between scenarios so that we maintain a constant γ-scale coverage (indicated by the slope of the tangential lines), the resulting pair of *β_C_* values become practically identical (compare with Fig. 2), which accurately reflects that both scenarios are equally aggregated at the patch scale. The advantage of standardizing γ-scale coverage becomes particularly clear when we consider the entire scaling relationship of *β_C_*. If we quantify *β_C_* for every possible value of *n* and plot them against γ-scale expected coverage, the values from large and small species pool fall on approximately the same line and the species pool dependence vanishes (Fig. 3 C, compare with 2 C).

**Figure 3:**
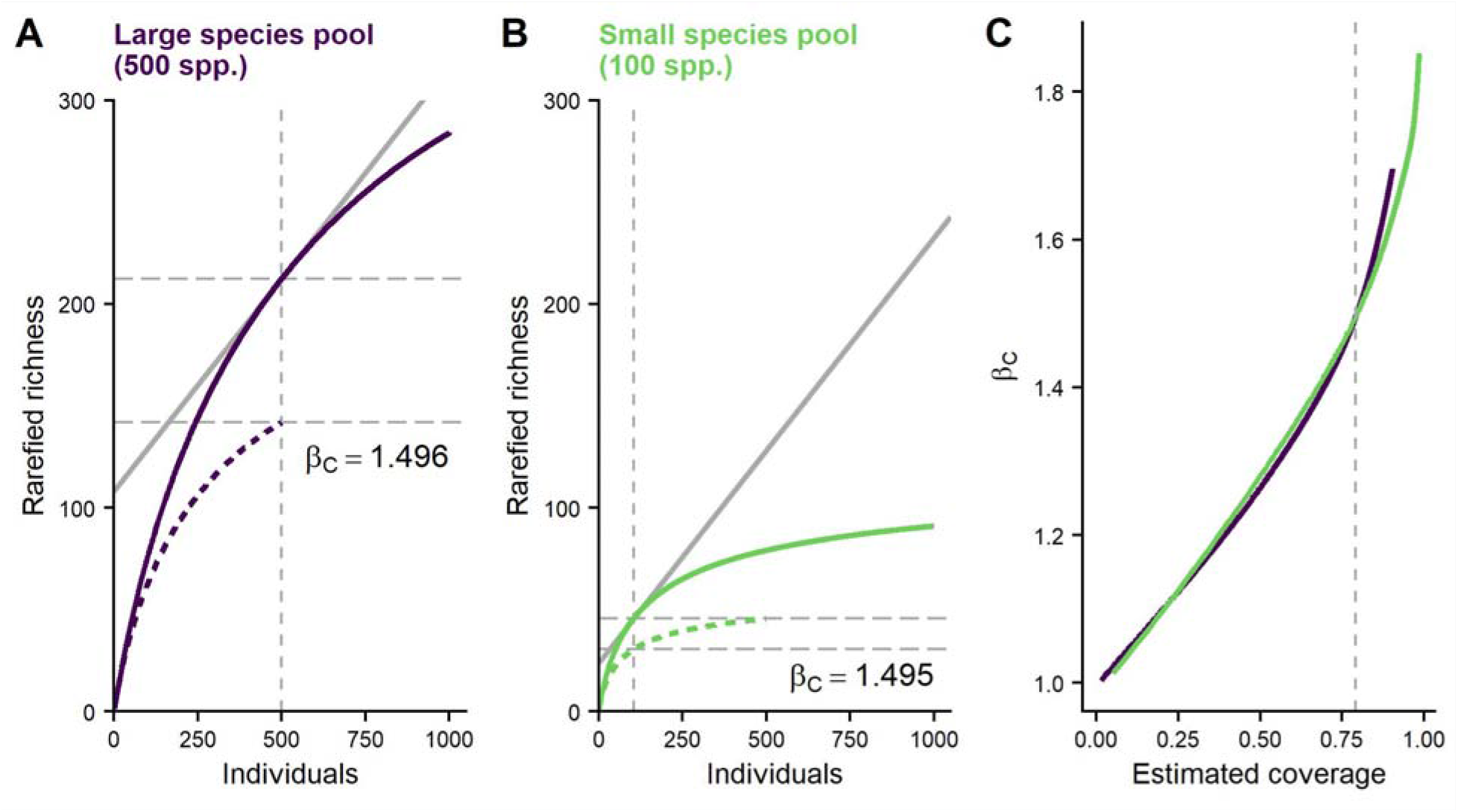
β_C_ is unaffected by species pool size in aggregated communities (compare with Fig 2). Two-scale IBRE curves for (A) a large species pool and (B) a small species pool. Solid curve: γ scale; dashed curve: α scale; dashed vertical grey line: number of individuals used for the calculation of β_S_n__ (i.e. the ratio of the horizontal dashed lines). Grey solid line shows the slope of the γ scale IBRE that relates to sample coverage (C=0.79 in both panels) C) β_C_ plotted as a function of expected coverage calculated at the γ scale. Dashed line marks the coverage value of 0.79 used in other panels.

To compare *β_C_* across multiple assemblages (e.g., with different species pools), we suggest the following protocol that makes use of interpolation and extrapolation.

1. Determine the appropriate target coverage value *C_target_* for the standardization:
  1.1. For each assemblage *j*, determine the smallest number of individuals observed at the α scale and call it *N_min_j__*.
  1.2. Using Eqn 4, estimate the expected γ-scale coverage *C_n_* corresponding to *N_min_j__* individuals, or up to *2N_min_j__* individuals if you wish to use extrapolation (Chao et al, 2014).
  1.3. Let *C_target_* be the smallest of the *C_n_* values across all assemblages.
2. Calculate *β_C_*:
  2.1. For each assemblage use the inverse of Eqn 4 to estimate the sample size *n_j_* corresponding to a γ-scale coverage of *C_target_*. Exclude all assemblages for which *n_j_* ≤ 1.
  2.2. Standardize γ and α scale species richness to *n_j_* individuals (using IBRE) to get *^γ^S_n,j_* and 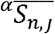.
  2.3. Calculate *β_C_* as:

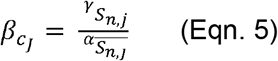

We implemented this procedure in an R package available at https://github.com/t-engel/betaC. It provides the function “C_target” that can be used for steps 1.1 and 1.2, and the function “beta_C” that carries out step 2.

This approach requires spatially replicated samples with abundance data (i.e., site by species matrices with abundance data), so that one can define at least two nested sampling scales (α and γ). We assume that the sampling design is standardized across all assemblages. This means there should be a consistent number of samples per assemblage and every sample should have the same effort (e.g., plot size, trap nights, etc.). Furthermore, we assume a consistent spatial extent at the γ-scale. If the number of samples, or their spatial extent, changes from one assemblage to the next, users should take a spatially constrained subset of the samples to keep extents as consistent as possible.

Like most measures of community differentiation, *β_C_* does not have an analytical variance estimator because there are no replicates at the γ-scale. Nevertheless, such variance is often desired, for example, when comparing spatial structure among different regions. To do so, we recommend calculating a distribution of *β_C_* for repeated random subsets of the samples. For example, one could use a Jackknife approach (i.e., systematically leaving out one sample at a time), use pairwise comparisons of samples, or comparisons among larger subsets of samples. While such variance can help to contextualize the observed values of β-diversity, caution should be taken because these calculations incorporate some degree of non-independence. Nevertheless, we provide an R function that carries out such resampling procedures for any number of samples (i.e. “betaC::beta_stand”).

### Proof of concept using simulations

To test the properties of *β_C_*, we simulated spatially explicit assemblages that varied in the size of the species pool and the degree of intraspecific aggregation using multivariate spatial point patterns. We used the R package mobsim (May et al., 2018) to carry out the simulations. Each simulated assemblage had 4000 individuals drawn from a lognormal SAD that was parameterized with a given species pool size, and a coefficient of variation equal to one. Then, we used the Thomas cluster process to distribute individuals in space, varying the degree of intraspecific spatial aggregation through the parameter that determines the number of conspecific clusters. The simulation was parameterized in a full-factorial design, where the species pool size encompassed every integer between 10 and 500, and the number of clusters was set to 1 (i.e., extreme intraspecific aggregation), 4, 10 and 20. To include a level that had no within species aggregation, we also implemented a random Poisson process to simulate completely random spatial distributions for species. Each combination of species pool and aggregation was replicated 3 times yielding a total of 7365 simulated communities. To sample from the regional communities, we placed 4 sample quadrats into each simulated community (Fig. 4 A, B). We calculated Whittaker’s 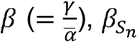, and *β_C_* among the 4 samples and examined their response to changes in species pool size and aggregation. Following the protocol above, *C_target_* was set to 0.55 and the sample size for *β_S_n__* was 50 individuals. The R code for the simulation is available on github (https://github.com/t-engel/betaC) and will be archived upon acceptance.

**Figure 4:**
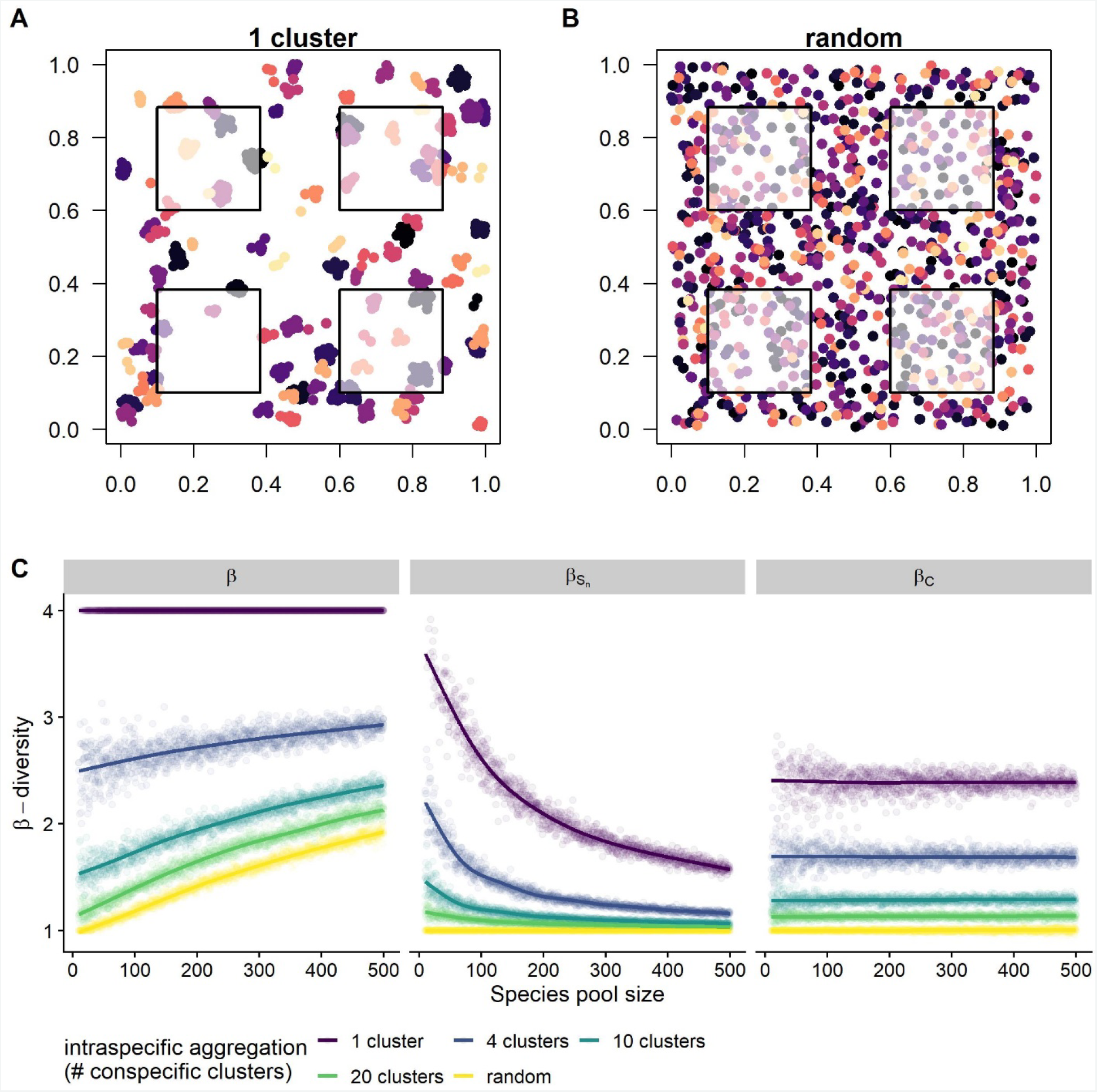
Simulated assemblages and the response of Whittaker’s β, β_S_n__ and β_C_ to changes in aggregation and species pool size. (A) Assemblage with extreme within species aggregation (1 cluster per species) and (B) assemblage where species have random spatial distribution. Species pool, SAD and numbers of individuals are constant between (A) and (B). Squares represent sample quadrats. (C) β-diversity metrics and their response to species pool size and intraspecific aggregation. Dots show the data. Lines show GAM fit to each metric with species pool size as the predictor; the GAM estimated separate smoothers for each level of intraspecific aggregation.

All three indices (Whittaker’s *β, β_S_n__, and β_C_*) were influenced by intraspecific community aggregation. Additionally, Whittaker’s *β* and *β_S_n__* were affected by the changes in species pool size, whereas *β_C_* was insensitive to this parameter. While both Whittaker’s *β* and *β_S_n__* responded to the degree of spatial aggregation and the species pool, they did so in contrasting ways, which is consistent with theoretical expectations (Fig. 4 C). For Whittaker’s *β*, the effect of the species pool decreased with increasing aggregation. This reflects that for strongly aggregated species distributions, the samples will always show high turnover regardless of the species pool. In contrast, under random species distributions, “spurious” (i.e. SAD related) sample differentiation is more likely to occur when there are many rare species that only occur in some of the samples (i.e. in large species pools). For *β_S_n__*, the effect of the species pool increased with increasing aggregation; as long as species are randomly distributed, *β_S_n__* is always one because the α IBRE curve falls onto the γ scale. However, when there is a deviation between the curves (as a result of aggregation), its magnitude for a given number of individuals (n) depends on the shape of the IBRE curves which in turn depend on the species pool (Fig 3). Only *β_C_* captures the spatial structure of the simulated communities independently of the species pool size because, by incorporating sample coverage, it adjust for the species pool-dependence.

We examined the robustness of *β_C_* using an alternative SAD (log-series), and by simulating spatial aggregation using the mean displacement length of the Thomas process (supplementary material S1). The results were qualitatively similar: *β_C_*, but not Whittaker’s *β* or *β_S_n__* responded to the changes of aggregation independently to changes of the SAD parameter (i.e., Fisher’s α). Additionally, we applied the null model by Kraft et al. (2011) to the simulated data and found that the measure of β-deviation, like *β_C_*, responded to the aggregation, but not to the species pool. Spearmans rank correlation between *β_C_* and β-deviation was 97.7 % which suggests that both approaches are measuring the same effect. (supplementary material S2).

### Empirical case studies

Next, we applied our approach to two forest datasets with varying species pool sizes. First, we used the Gentry Forest plot dataset (Gentry, 1988; Phillips & Miller, 2002), which has been used to address the question of how spatial aggregation varies with latitude (Kraft et al. 2011), and has frequently been the subject of a debate on how to formulate appropriate null models for β-diversity (Qian & Song, 2012; Qian et al., 2013; Xu et al., 2015; Ulrich et al., 2017). We computed Whittaker’s *β* and *β_C_* among the subplots of the sites located in the Americas. As expected from the difference in species pool size along this gradient, and shown by previous studies (e.g., Kraft et al., 2011), Whittaker’s β declined with latitude (Fig. 5A). In contrast, *β_C_*, which controls for species pool related sampling effects, showed no significant change along the latitudinal gradient (Fig. 5B). Given this, we conclude that at the given scale, there is no evidence for a change of spatial aggregation along this gradient. Declines in Whittaker’s β with increasing latitude appears to be mostly driven by changes in the size of the regional species pool rather than changes in the spatial structuring of individuals. Importantly, although we come to qualitatively similar results as the abundance-based null models of Kraft et al. (2011) and Xu et al. (2015), our method has the advantage of explicitly incorporating an estimate of sample completeness. Rather than simply shuffling site-by-species matrices, users are confronted with the completeness of their samples as part of the analytical workflow, and prior to interpreting any results. In this case, our analysis provides quantitative evidence for the argument that these types of forest plots may be too small to robustly compare spatial patterns of diversity and any associated differences in community assembly, especially in the tropics (Tuomisto & Ruokolainen, 2012). *C_target_* is set by the site with the lowest coverage (C_n_), which for the Gentry Forest Plot data was *C_target_* = 0.09. This means that inferences are being drawn from a sample of only approximately 9 percent of the individuals in the assemblage.

**Figure 5:**
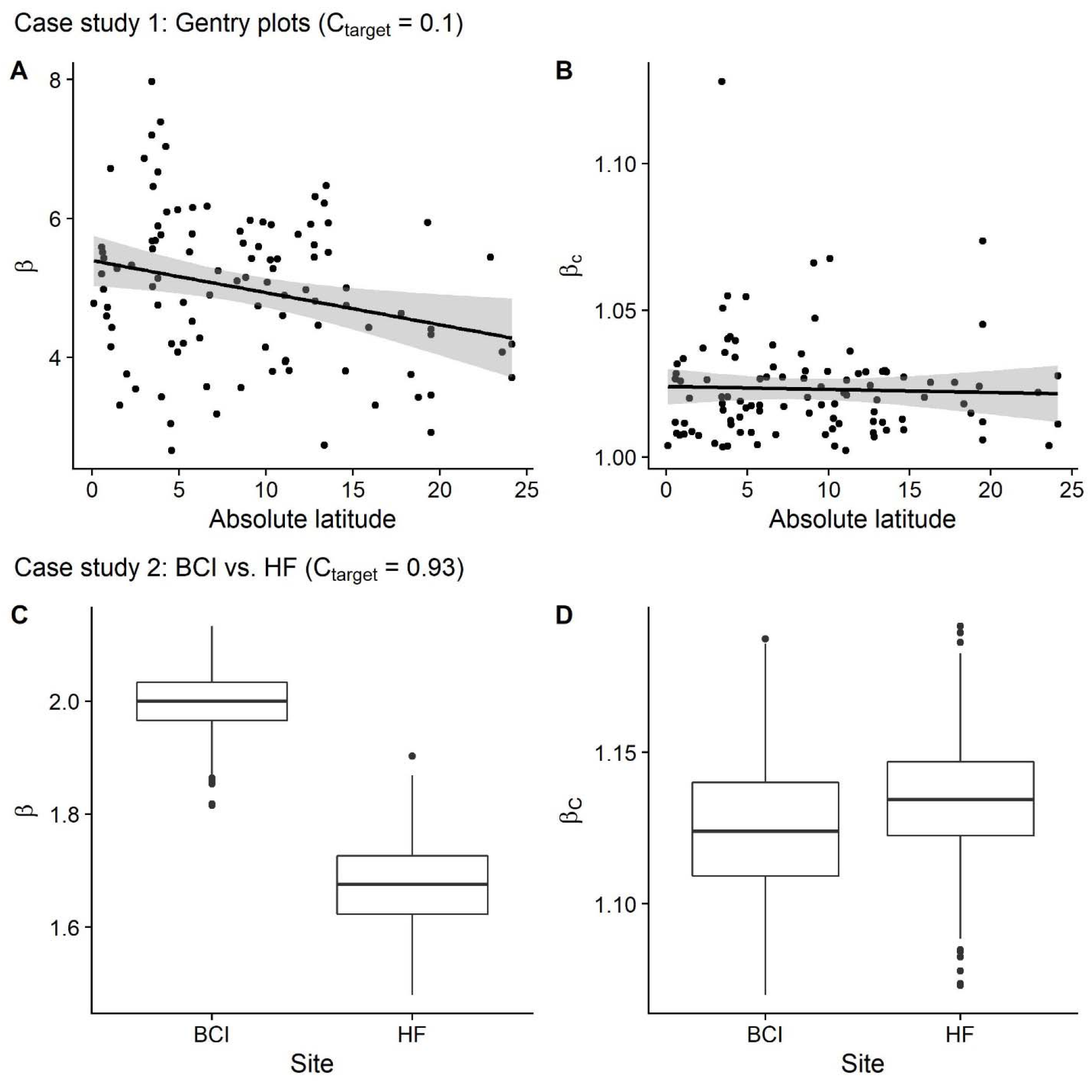
Case studies exploring β-diversity along (A, B) a latitudinal gradient of Gentry forest plots and (C, B) comparing Barro Colorado Island (BCI) and Harvard Forest (HF). Both examples show significant changes in Whittaker’s β (A,C) while β_C_ (i.e. β standardized for sample coverage) showed no significant change. Solid lines show simple linear regressions. Boxplots show distribution of values from 1000 redraws of 10 samples, respectively. Whiskers: non-outlier range; box: interquartile range: bar: median

For a second case study, we explored a similar question along the latitudinal gradient, but with larger plots, so that a more substantial fraction of the species pool would be sampled. To do so, we compare the spatial structuring of trees within the temperate forest plot at Harvard Forest (HF), in the northeastern U.S. (Orwig et al, 2015), with the well-studied tropical rainforest plot at Barro Colorado Island (BCI), Panama (Condit et al, 2019). Because the data from HF was from only 35 hectares, we took a 35 ha subsection of the 50 ha plot at BCI, and found a considerable difference in γ-diversity; 38 species were present at HF and 217 species were in the analyzed section of BCI. For both locations, we used our resampling approach to calculate a distribution of β-diversity at the scale of ten 1 ha subplots (using 1000 random draws with replacement). Again, we found that Whittaker’s β was consistently higher in the more diverse site (BCI). While HF had an expected α-scale coverage of 99%, it was 90% at BCI. By Interpolating HF and extrapolating BCI (following the protocol above), we standardized both sites to a target coverage of 0.93, and found no meaningful difference between in the corresponding *β_C_* (Fig.5 D). In short, as with our analyses of the Gentry data above, but with samples that more adequately characterise the assemblages of interest, we find that the observed differences in Whittaker’s β from temperate to tropical forests were largely expected given the differences in the species pool. That is, once species pool related sampling effects were taken into account, there do not appear to be any meaningful differences in the spatial structuring of these two forests.

## Discussion and conclusions

Building on previous work using rarefaction and coverage-based approaches (e.g., Colwell and Coddington, 1994; Chao and Jost, 2012; Chao et al., 2014; Chase et al., 2018; McGlinn et al., 2019), we developed a metric standardized by sample coverage to quantify the degree of intraspecific spatial aggregation, independent of changes in the size of the species pool and the regional SAD. Our theoretical considerations and simulations of spatially explicit assemblages show that *β_C_*, remains unaffected by changes in the species pool, which allows for comparisons of intraspecific aggregation along large biogeographic gradients. Our empirical case studies suggest that the magnitude of intraspecific aggregation does not change along a latitudinal gradient of forest plots. Importantly, our method requires analysts to determine the target completeness using information contained in the samples from their study. In the case of the commonly used Gentry plots, this shows that the samples cover only a small fraction (~10%) of the individuals in the underlying assemblages, and may therefore of limited use for making inferences about their small-scale spatial structure (see also Tuomisto & Ruokolainen, 2012).

Our approach represents an important advance over existing methods to measuring spatial aggregation because of its strong link to existing biodiversity sampling theory. Specifically, we use the γ-scale rarefaction curve as an analytical null model for the expected α-diversity in the absence of spatial structure. Whilst conceptually similar to existing null models (e.g. Kraft et al, 2011), our approach has several advantages. First, it bypasses the computation of Whitaker’s beta, and subsequently β-deviation. Instead, we measure the deviation between γ and α scale directly from the IBRE curves. *β_C_* can be thought of as the factor by which spatial structure has reduced α diversity compared to the random expectation. This makes it more intuitive than β-deviation (e.g., Kraft et al. 2011, Xing & He, 2020) because it can be directly interpreted as an effective number of distinct communities (Jost, 2007), conditional on the estimated sample coverage. Additionally, our approach explicitly incorporates an estimate of sample completeness into the analytical workflow.

Ulrich et al. (2017) have argued that null model approaches are limited in their ability to disentangle species pool and aggregation effects, unless they incorporate external data on the sizes of the relevant species pools. Here, we make use of the idea that the sample itself can also provide an estimate of its completeness (Good, 1953; Chao and Jost 2012). Our method uses the shape of the IBRE curve, itself determined by the SAD, to draw inferences for an estimated constant fraction of the individuals in the underlying community (i.e., a constant sample coverage). This approach to standardization implicitly assumes that there is an asymptote in species diversity (Chao and Jost, 2012). While this assumption is mathematically convenient, it cannot strictly be true; due to species aggregation at higher scales we will always find more species with more samples, until the entire global pool is sampled (Williamson et al. 2001). Nevertheless, *β_C_* does not extend to the asymptote itself, but merely employs a useful approximation of sample completeness via sample coverage. As spatial structure is an inherently scale-dependent phenomenon, with this approach we can only measure it at the spatial scale prescribed by the samples (i.e. spatial structure within the γ scale).

While *β_C_* isolates the degree of spatial aggregation, it does not have some of the properties that are sometimes considered essential for β-diversity metrics (Tuomisto 2010 a; Jost, 2007. For example, traditional β-diversities range between unity and the total number of sampling units, and they can be transformed into N-community (dis)-similarities in the range [0,1] (Jost 2007; Chao and Chiu, 2016). In contrast, *β_C_* can only reach the number of samples when the samples have a coverage of 1 (i.e. *C_target_* is 100%) (see supplementary material S3). In such cases, where the curves have reached an asymptote, *β_C_* equals Whittaker’s *β* and one can derive the corresponding (dis-)similarity. However, for incomplete samples, the N-community transformation of β-diversity is generally not recommended (Chao and Chiu, 2016), and we consider it a strength of our method that it exposes such situations. Only if *C_target_* is 100%, pairwise (dis)-similarities will not be affected by the species pool.

Although our approach accounts for differences in sample completeness, it still requires standardized sampling (or data that could be standardized *post hoc*) to make valid inferences. The sampling effects we treat here arise passively due to differences in species pools, and not as a result of different sampling strategies and/or effort. For example, forest plots in our case studies were completely sampled only in the sense that every tree was counted in a given area or subplot. However, with respect to the regional species pool or even just the observed γ diversity, a subplot of a given size in the tropics is likely a much less complete sample, compared to an equally-sized subplot a temperate region, even when all individuals are counted in each subplot. It is this interaction of sampling effort and the size of the species pool that leads to the null expectation of increasing sample differentiation with increasing species pool size, and that our method adjusts for.

In conclusion, our approach allows us to explicitly disentangle non-random spatial patterns of species diversity (e.g., intraspecific aggregation) amidst variation in species pool size and associated sampling effects. In conjunction with other diversity metrics sensitive to diversity components such as the SAD and total community abundance (see Chase et al., 2018; McGlinn et al., 2019), *β_C_* allows deeper insights into how spatial structuring within communities influences patterns of biodiversity and its change. Applications could, for example, shed light onto the assembly processes that govern (meta-)communities along biogeographic gradients, and contribute to a better understanding of the spatial diversity patterns that underlie the scale-dependent biodiversity trends observed during the current biodiversity crisis.

## Acknowledgements

We acknowledge the support of the German Centre for Integrative Biodiversity Research (iDiv) Halle-Jena-Leipzig funded by the German Research Foundation (FZT 118). In addition, we thank Xiao Xiao, Tiffany Knight, Leana Gooriah, Petr Keil and Ingmar Staude for discussions and feedback on the approach.

