## supplementary material for "Using coverage-based rarefaction to infer non-random species distributions"

### S1. Additional simulation

We carried out an additional simulation where we parameterized the species pool using the log-series distribution and variation in the alpha parameter. Additionally, we simulated different degrees of spatial aggregation by changing the mean displacement length (sigma) of the Thomas process (as opposed to the number of clusters as presented in the main text). We used all integers from 1 to 100 as alpha values for the log-series SAD which resulted into species pools of 5 to 344 species. We used the following sigma parameters to get increasingly aggregated species distributions: 1, 0.8, 0.4, 0.2, 0.1, 0.01. The number of clusters was set to 1 for each species. Like in the analysis presented in the main text, we found that Whittaker’s beta and beta_Sn responded to the change in SAD (alpha parameter), while $\beta_{C}$remained mostly unaffected by it (Fig S1).

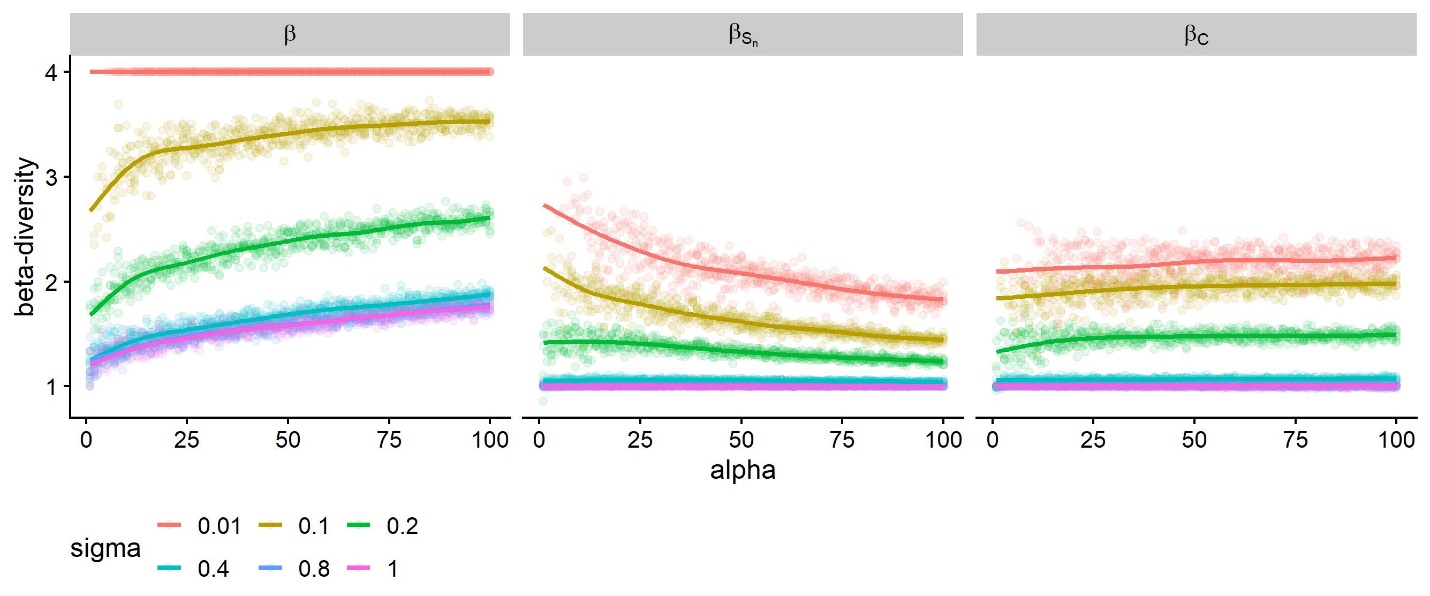

*Fig S1: Response of Whittaker’s* $\beta$*,* $\beta_{S_{n}}$ *and* $\beta_{C}$ *to changes in aggregation and species pool size from the additional simulation*

### S2. Beta-deviation

We also applied the Kraft null model to the simulated data. We used the code provided by Sebastian Tello provided on his website: <http://jsebastiantello.weebly.com/r-code.html>. We reshuffled the simulated site-by species abundance matrices 400 times, keeping the SAD and number of individuals per site constant. For each permutation, we calculated Whittakers beta. Then we calculated beta-deviation as:

$$\beta_{dev}=\frac{\beta_{obs}-mean(\beta_{null})}{sd(\beta_{null})}$$

Where $\beta_{obs}$ is the observed beta-diversity, $mean(\beta_{null})$ is the mean of the null distribution and $sd(\beta_{null})$ is the standard deviation.

Figure S2 shows how beta-deviation responds to our simulation parameters. Compared to $\beta_{C}$, it still shows some week species pool dependence for intermediate levels of intraspecific spatial aggregation.

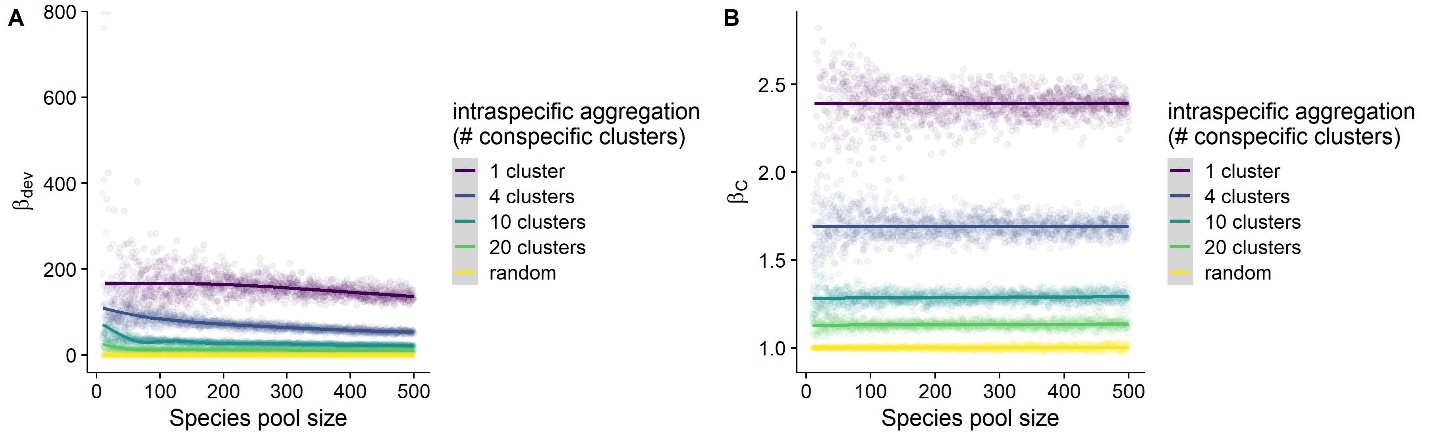

*Fig S2: Response of (A) beta-deviation and (B)* $\beta_{C}$ *to simulation parameters.*

### S3. Asymptotic behavior of $\beta_{C}$

Many authors have argued that metrics of beta-diversity should range between 0 and the number of sampling units (e.g. Jost, 2007). $\beta_{C}$ shows this behavior asymptotically. In the main text, we illustrate our method using an example with communities from differently sized species pools (Fig 2 and Fig 3). Although in both cases turnover is assumed to be at a maximum, the value of $\beta_{C}$ does not reach the number of sampling units 2. This is because in this example the samples are not complete with respect to the total species pool (C_target= 0.8). $\beta_{C}$ does range between 0 and the number of sampling units if target coverage is 100%. In this case $\beta_{C}$ is exactly the same as Whittaker’s beta, as both alpha and gamma scale curve have reached an asymptote that corresponds to the observed species richness.

Fig. S3 shows the same example but with an increased number of individuals sampled from each of the patches. Here, $\beta_{C}$ can be calculated for a coverage of 1 (i.e. vertical dashed line) and its value becomes 2. When the value of is smaller for lower coverages, this merely reflects the fact that a fraction of the individuals in the assemblage are not covered by the species in the samples.

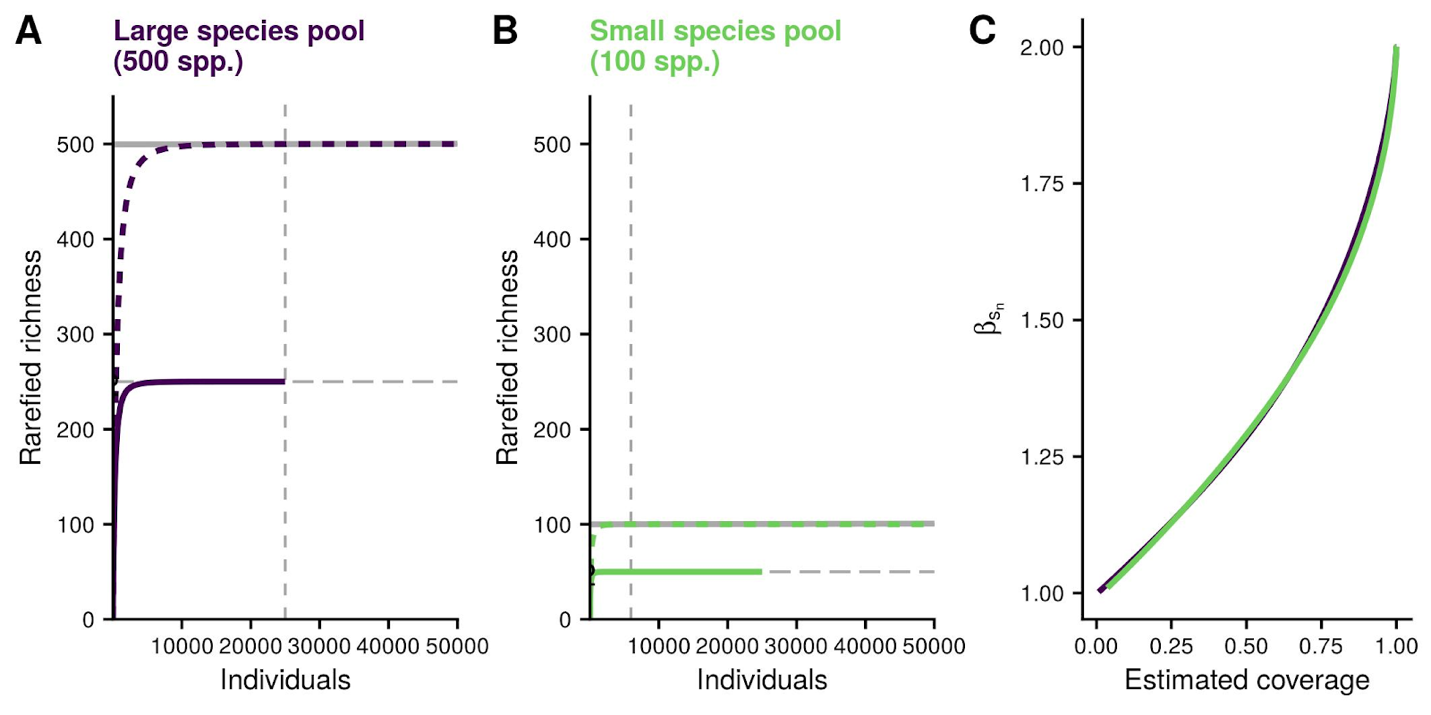

*Fig S3: Under a scenario of complete turnover,* $\beta_{C}$ *reaches the number of sampling units (here 2) when sample coverage is 100%*
